# Human NHE1 binds full-length calcineurin with high affinity and inhibits substrate dephosphorylation

**DOI:** 10.64898/2026.09.10.750614

**Authors:** A. Sorg, J.-S. Haas, M. Heinrich, S. Gauss, B.A. Kern, V. Timmermann, E.V. Mymrikov, C. Hunte

**Author notes:** contributed equally (A. Sorg, J.-S. Haas). correspondence to: Institute of Biochemistry and Molecular Biology, ZBMZ, Faculty of Medicine, CIBSS, University of Freiburg, Stefan-Meier-Strasse 17, 79104 Freiburg, Germany; (EVM), (CH).

## Abstract

Calcineurin (Cn) interacts with the Na^+^/H^+^-exchanger NHE1 through its calcineurin-binding domain (NHE1-CnBD). How activation of full-length Cn influences this interaction and its functional consequences has remained unclear. Here, we quantified the interaction between full-length human Cn and NHE1-CnBD in distinct activation states using fluorescence anisotropy and quantitative size-exclusion chromatography. Ca^2+^/calmodulin activation increased Cn affinity for NHE1-CnBD from submicromolar to tens of nanomolar affinity and stabilized the complex. NHE1-CnBD outcompeted the immunosuppressive cyclosporin A/cyclophilin A complex for Cn binding and selectively inhibited dephosphorylation of a phosphopeptide substrate while enhancing hydrolysis of *p*-nitrophenyl phosphate. These findings identify NHE1 as a high-affinity, substrate-competitive Cn regulator that blocks substrate recognition rather than catalysis, providing a mechanistic framework for reciprocal regulation between Cn and NHE1.

## INTRODUCTION

As the major ubiquitous proton extruder at the plasma membrane, the sodium/proton exchanger 1 (SLC9A1, NHE1) is central to intracellular pH and cell volume homeostasis and thereby supports diverse cellular processes including proliferation, migration, and survival [1]. NHE1 consists of a membrane transport domain followed by a long largely intrinsically disordered cytoplasmic regulatory region (NHE1-CTD) [2]. NHE1-CTD serves as a regulatory hub for several interacting proteins that modulate NHE1 activity [1], including the Ca^2+^-binding proteins calcineurin B homologous proteins (CHPs) [3, 4], calmodulin (CaM) [5, 6] and calcineurin (Cn) [7, 8].

Cn is a heterodimeric Ca^2+/^CaM-dependent Ser/Thr phosphatase that plays a central role in immune responses, cardiac hypertrophy, neuronal signalling, and other Ca^2+^-dependent signalling pathways [9, 10]. It recognizes the substrates primarily through two short linear motifs (SLiMs), termed PxIxIT and LxVP after the consensus amino acid sequences first identified in NFAT (nuclear factor of activated T-cell) transcription factors [11–13]. Under resting intracellular Ca^2+^ concentrations (∼100 nM), Cn is maintained in an autoinhibited state (Fig. 1A). Binding of Ca^2+^ to the regulatory B subunit partially activates the enzyme, whereas full activation requires binding of Ca^2+^-loaded calmodulin (CaM) to the catalytic A subunit, which results in displacement of the autoinhibitory domain (AID) from the catalytic site [14, 15]. In addition to stimulating phosphatase activity, CaM binding enhances the interaction of Cn with NFAT1 and Ca_V_3.2 (T-type voltage-gated Ca^2+^ channel)-derived peptides [16, 17], suggesting that activation also regulates substrate recognition. However, the influence of activation on target-binding affinity has not been quantified for full-length heterodimeric Cn.

**Fig. 1.**
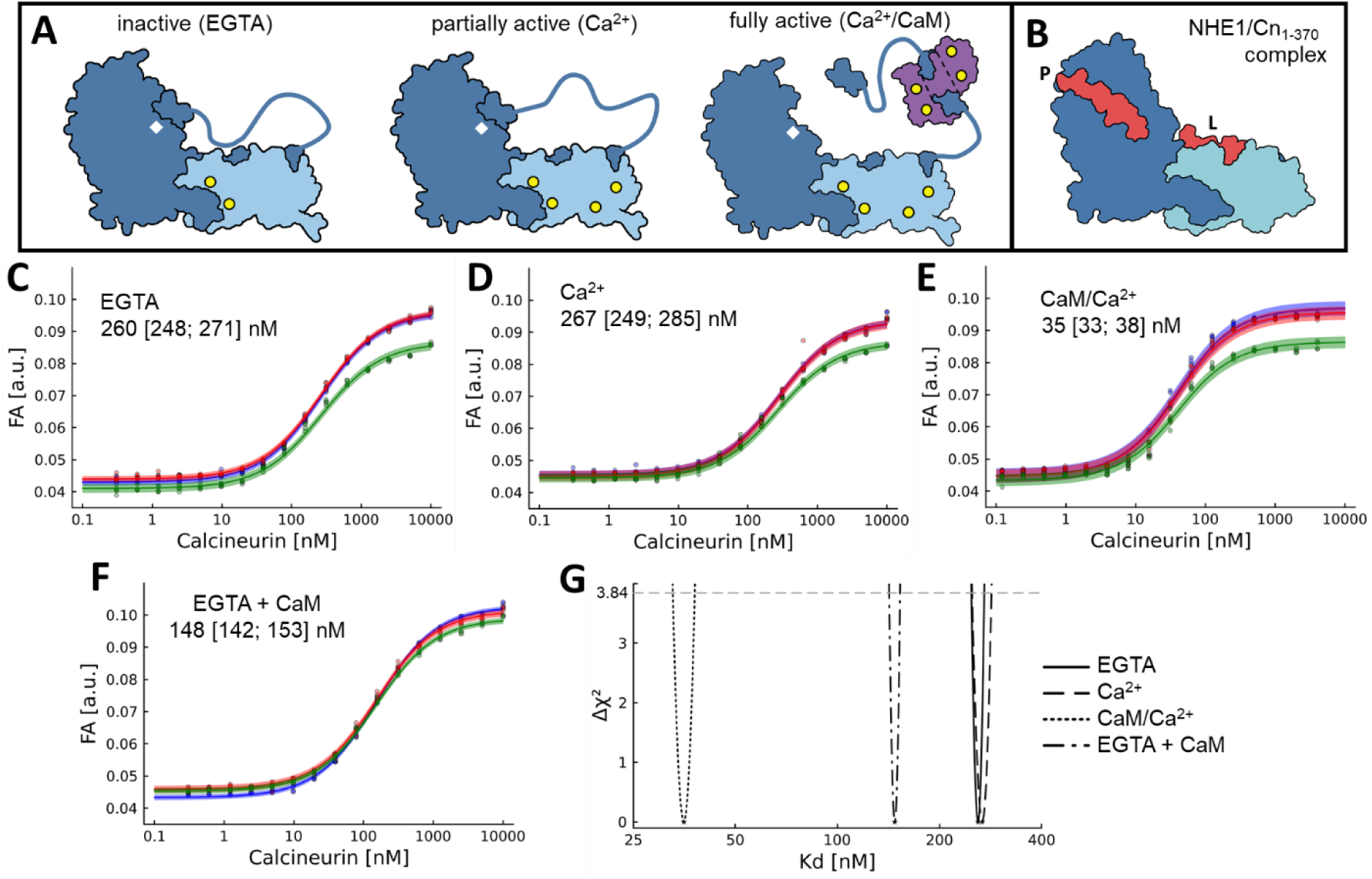
Ca^2+^/CaM activation of full-length calcineurin significantly increases its affinity for NHE1-CnBD. **A**, Schematic representation of the three calcineurin (Cn) states. Inactive Cn: CnA (dark blue), CnB (light blue) with two Ca^2+^ ions (yellow circles), CnA autoinhibitory sequence (AIS) of regulatory region (dark blue line) interacts with LxVP-binding site, CnA C-terminal autoinhibitory domain (AID) blocks the catalytic site (white diamond). Partially active Cn: CnB with two additional Ca^2+^ ions, AIS detached. Fully active Cn: Ca^2+^-loaded CaM (purple) bound to CnA regulatory region, AID released. Scheme based on structures of Cn (PDB ID: 1AUI [20]) and the CaM-bound regulatory region of Cn (PDB ID: 4Q5U [33]). **B**, Schematic representation of complex of NHE1-CnBD (red) and constitutively active Cn_1-370_ (PDB ID: 6NUC [7]). The PIVITID motif (P) binds to the PxIxIT-binding site, whereas the LTVP motif (L) binds to the LxVP-binding site. Ca^2+^ ions bound to CnB are omitted for clarity. **C-F**, Binding of fluorescein-labelled NHE1-CnBD_Cys_ (NHE1-CnBD*) to Cn_WT_ measured by fluorescence anisotropy. **C**, inactive Cn, 5 mM EGTA, **D**, partially active Cn, 1 mM Ca^2+^, and **E**, fully active Cn, 1 mM Ca^2+^ and 8 µM CaM). **F**, inactive Cn in presence of CaM (5 mM EGTA, 10 µM CaM). Measurements were performed with three independent biological replicates (in different colour) and 4 (**C-E**) or 3 (**F**) technical replicates each; The combined data were fitted with a 1:1 binding model using global non-linear regression. Derived *K*_D_ values and 95 % confidence intervals are included as insets. **G**, Comparison of 95 % confidence intervals of *K*_D_ values from **C-F** calculated with profile likelihood method.

The interaction between NHE1 and Cn appears to regulate the activities of both binding partners [7,8,13]. Cn dephosphorylates Thr779 located downstream of the binding sites, suggesting direct regulation of NHE1 [7]. Conversely, NHE1-mediated local intracellular alkalinization promotes Cn/NFAT signalling in cardiomyocytes [8], whereas disruption of Cn signalling alters NHE1 regulation in neurons [18]. These observations indicate reciprocal functional coupling between NHE1 and Cn. Given the ubiquitous presence of both proteins, this interplay may contribute to Ca^2+^-dependent signalling in multiple tissues [1,13].

Cn is also a major therapeutic target of the immunosuppressants cyclosporin A (CsA) and FK506 (tacrolismus). In complexes with their respective immunophilins, cyclophilin A (CypA) and FKBP12, these drugs occupy the LxVP-binding site of Cn and prevent substrate-recognition without directly interfering with the catalytic site [19, 20]. Thus, understanding how endogenous proteins bind to this regulatory surface is important both for defining the molecular basis of substrate recognition and for understanding how pharmacological inhibition may perturb native signalling complexes.

Cn interactions span a remarkably broad affinity range. Most physiological substrates bind with micromolar affinity, allowing rapid substrate turnover [15, 21], whereas endogenous inhibitors and regulators such as RCAN1 and CABIN1 bind with submicromolar to low-nanomolar affinity through multivalent interactions involving two or three binding sites [22–24]. Notably, most quantitative studies have employed isolated CnA or constitutively active truncated variants lacking the AID and the CnA regulatory region, whereas the binding properties of full-length heterodimeric Cn remain poorly characterized. The regulatory region preceding the AID contains an autoinhibitory sequence (AIS) that resembles an LxVP motif and occupies the corresponding binding site [13, 25]. Consequently, measurements with constitutively active Cn may overestimate the affinity of physiological binding partners.

An LxVP motif (LTVP) and a PxIxIT-like motif (PIVITID) are located within the NHE1-CTD. They are separated by a flexible linker that enables simultaneous binding to both binding sites of Cn [7]. Studies using a constitutively active truncated Cn variant showed that both motifs are required for high-affinity binding and that their bivalent interaction enhances affinity through avidity [7]. As accessibility of the LxVP-binding site is thought to depend on Ca^2+/^CaM activation [14], it remains unknown how the activation state of full-length Cn influences NHE1 binding or whether bivalent NHE1 binding, in turn, modulates Cn phosphatase activity.

Here, we quantitatively characterize the interaction between the NHE1 Cn-binding domain (NHE1-CnBD) and full-length human Cn in inactive, partially active and fully active states. We show that Ca^2+^/CaM activation increases the affinity of Cn for NHE1-CnBD more than sevenfold and markedly stabilizes the complex. NHE1-CnBD binds Cn more strongly than the immunosuppressive CsA/CypA complex and selectively inhibits dephosphorylation of a phosphopeptide substrate while promoting hydrolysis of a small-molecule substrate. These findings identify the NHE1 Cn-binding region as a high-affinity substrate-competitive regulator of Cn and provide a mechanistic framework for reciprocal regulation between these two signalling proteins.

## MATERIALS AND METHODS

### Plasmids and Proteins

*Escherichia coli*-optimized coding sequences of full-length human Cn catalytic (CnA, α isoform, UniProtKB Q08209) and regulatory (CnB, P63098) subunits cloned into pETDuet-1 vector (pETDuet-1_His6-CnA/CnB), and cyclophilin A (CypA, P62937) cloned into pET30a (pET30a_CypA), were obtained commercially (Biocat GmbH, Heidelberg, Germany).

The plasmid encoding constitutively active Cn (pETDuet-1_His6-CnA1-370/CnB), comprising CnA residues 1-370 and full-length CnB (hereafter Cn_1-370_) was generated by deleting the coding sequence for CnA residues 371-521) using NEBuilder^®^ HiFi DNA Assembly (New England Biolabs, Ipswich, MA, USA). The same cloning strategy was used to generate all other constructs. The coding sequence of untagged CaM (P0DP23) was cloned from pQE-T7_CAM1 [3] into pET30a(+) (pET30a_CAM1). His-MBP-CnBD (NHE1 residues 674-730, P19634) was generated by replacing the 3PDZ1^KSTQF^ insert in His-MBP-3PDZ1^KSTQF^ plasmid [26] with the corresponding NHE1 sequence amplified from pRSF13-SLC9A1^CTD^ [3] (pET30a_His8-MBP- CnBD). The S729C substitution was introduced during subcloning of the pET30a_His-MBP- CnBD_Cys_ plasmid using mutagenic primers to generate the NHE1-CnBD_Cys_ variant. The pET30a_His-GFP-CnBD plasmid was generated by replacing the URAT1 peptide sequence in the previously described GFP^URAT1^ plasmid [26] with the NHE1-CnBD sequence.

All proteins were produced in *E. coli* BL21(DE3) cells transformed with the corresponding plasmid and grown in LB medium supplemented with 50 µg/ml kanamycin for pET30a(+)- based plasmids or 100 µg/ml carbenicillin for pETDuet-1-based plasmids. Expression was induced at OD600 = 0.6 with 0.5 mM IPTG followed by incubation for 5 h at 30° for NHE1- CnBD constructs, and overnight at 20°C for Cn, at 30° for CaM, and at 37° for CypA.

All chromatography columns used in the study were from Cytiva (Marlborough, MA, USA). Full-length heterodimeric calcineurin (Cn_WT_) and its constitutively active variant, Cn_1-370_, were purified essentially as described previously [7, 27] without removal of the poly-histidine tag. Briefly, Cn_WT_ was purified by immobilized metal affinity chromatography (IMAC) using a HisTrap FF 5 ml column, followed by gel filtration using a Superdex200 HiLoad 26/600 column and as final step by ion-exchange chromatography (IEX) using a HiTrapQ 5 ml column. Cn_1-370_ was purified by IMAC, IEX and final gel filtration using the same columns.

NHE1-CnBD and CnBD_Cys_ were purified according to the protocol previously developed for the human PDZK1 PDZ1 domain [26], consisting of two IMAC steps separated by TEV-protease cleavage, and a final gel filtration using a Superdex75 HiLoad 26/600 column. The GFP^NHE1-CnBD^ fusion protein (GFP^NHE1-CnBD^) was purified as described previously for GFP^URAT1^ [26]. Recombinant untagged CaM was purified essentially as described previously [28, 29] by Ca^2+^-dependent hydrophobic interaction chromatography (HIC) on a HiTrap Phenyl HP column after heat clarification of the lysate at 70°C, followed by gel filtration using a Superdex75 HiLoad 26/600 column. CypA was purified according to Liu et al. [30].

### Fluorescence anisotropy

Binding affinities of Cn for NHE1-CnBD under different conditions were determined by fluorescence anisotropy (FA). NHE1-CnBD_Cys_ was labelled at 20 µM final concentration with a 20-fold molar excess of fluorescein-5-maleimide (Cayman Chemical, Ann Arbor, MI, USA) in FA buffer (20 mM HEPES, 150 mM NaCl, 1 mM CaCl_2_, 1 mM MgCl_2_, 1 mM TCEP, pH 7.5) for 30 min at room temperature in the dark.

Excess dye was removed by three rounds of dialysis against FA buffer with concentrating steps in between, followed by buffer exchange using a PD-10 column (Cytiva). About 70 % degree of labelling for NHE1-CnBD* (fluorescein-labelled NHE1-CnBD_Cys_) was achieved. Aliquots of NHE1-CnBD* in FA buffer were flash frozen in liquid nitrogen and stored at -80°C.

For affinity measurements, Cn in FA buffer was serially titrated (1:1 dilution, 16 steps) in black flat-bottom 384-well plates with 30 µl final volume using a Pipetting robot Liquid Handling Station (Brand GmbH + Co KG, Wertheim, Germany). An equal volume of a master mix containing 20 nM of NHE1-CnBD* and 0.2 % BSA (Sigma-Aldrich, St. Louis, MO, USA, #A7906) in FA buffer including supplements was added to each well. Binding to inactive Cn was measured in the presence of 5 mM EGTA, whereas fully active Cn was analysed in the presence of 8 µM CaM. Binding was also measured in the presence of both 5 mM EGTA and 10 µM CaM. To assess the effects of Cn inhibitors, the assay was performed in the presence of 15 µM CsA/CypA with or without 10 µM CaM. As a control, 15 µM CypA alone was included. After incubation for 30 min at room temperature in the dark, parallel and perpendicular polarized fluorescence intensities were measured at 25°C using a Clariostar Plus plate reader (BMG Labtech, Ortenberg, Germany) with excitation and emission filters of 482/16 nm and 530/40 nm, respectively. Detector gain for both polarization channels was adjusted to yield a fluorescence polarization of 75 mP for the NHE1-CnBD*-only control. Background fluorescence was determined from samples lacking NHE1-CnBD* and subtracted before FA values were calculated using MARS software (BMG Labtech).

Measurements were performed using three independent biological replicates of Cn, each analyzed with three to four technical replicates. To determine *K*_D_ values, the global fit analysis using a 1:1 binding model [26] was performed as described earlier [31] using the Julia package InformationGeometry.jl [32]. The *K*_D_ parameter represents a biological quantity, that is global and shared between the biological replicates, while the remaining parameters (r_max_ and r_0_) are affected by experimental procedure and therefore different between biological replicates. Additionally, for each biological replicate, the dose-independent measurement noise was estimated.

### Analytical gel filtration

Quantitative analytical gel filtration was performed as described previously [26] using a Superdex 200 Increase 10/300 GL column equilibrated with FA buffer. Protein elution was monitored by absorbance at 280 nm (A_280_) for total protein and at 490 nm (A_490_) for GFP^NHE1-CnBD^. Individual proteins were loaded at 10 µM (alone and in mixtures) except CsA and CypA, which were used at 15 µM each). After 30 min incubation, a 500 µl sample was injected using an ALIAS autosampler (Cytiva). Individual protein concentrations along the elution profile were quantified from A_280_ and A_490_ as described previously for PDZK1/URAT1 complexes [26] using equations 1-2.

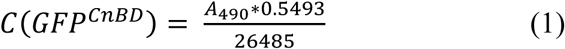

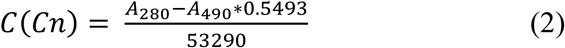

For the sample, containing Cn, CaM and GFP^NHE1-CnBD^, the concentration of the Cn/CaM complex was calculated according to equation 3 assuming the 1:1 stoichiometry due to a picomolar affinity between Cn and CaM in the presence of Ca^2+^ [14, 33, 34].

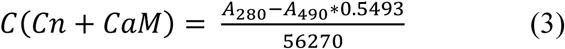

SEC analysis of the CsA/CypA effect on the NHE1-CnBD–Cn complexes was done following the same protocol, but without quantification of the protein concentrations.

### Calcineurin phosphatase assay

Phosphatase activity was measured using either the small-molecule substrate *para*-nitrophenyl phosphate (pNPP) or the Cn-specific RII phosphopeptide substrate. All reactions were performed in 96-well clear-bottom plates.

#### pNPP assay

The reaction mixture contained 66 mM pNPP in activity buffer (20 mM Tris-HCl, 100 mM NaCl, 1 mM CaCl_2_, 1 mM MgCl_2_, 0.5 mM TCEP, 0.05 % NP-40, pH 7.5). Depending on the experimental condition, the buffer was used as prepared (partially active Cn_WT_ or Cn_1-370_), supplemented with 5 mM EGTA (inactive Cn_WT_), or with 100 nM CaM (fully active Cn_WT_). Reactions were initiated by addition of Cn to the reaction mix. Absorbance at 410 nm was recorded every 2 min for 40 min (Cn_WT_) or every minute for 20 min (Cn_1-370_) at 30°C using a PowerWave XS2 plate reader (BioTek, Winooski, VT, USA). Phosphatase activity was calculated from the linear part of the curves (2-20 min) using ε_410_ = 14 000 M^-1^cm^-1^ for *para*- nitrophenol at pH 7.5 [35]. Activity measurements in the presence of 50 nM, 100 nM, or 1.0 µM of NHE1-CnBD peptide were performed for Cn_WT_ and Cn_1-370_, each with three technical replicates.

#### RII phosphopeptide assay

Reactions were initiated by addition of RII phosphopeptide (MedChemExpress, Monmouth Junction, NJ, USA) at 0.15 mM final concentration (in activity buffer) to the reaction mixture which contained 10 nM Cn_WT_ and 100 nM CaM either with or without 50 nM, 100 nM, or 1.0 µM of NHE1-CnBD peptide (in activity buffer). Activity of partially active Cn_WT_ was measured without CaM and NHE1-CnBD addition. Activity of Cn_1- 370_ was measured the same way in the absence of CaM. Reactions were performed in triplicates in the final volume of 50 µl at 23°C and stopped after 0-60 min by addition of 100 µl BIOMOL Green (Enzo Life Sciences, Lörrach, Germany). Absorbance at 620 nm (A_620_) was measured with the same plate reader. A calibration curve of 0-50 µM phosphate standard (Enzo Life Sciences) was used to convert A_620_ into phosphate concentrations.

For both assays, phosphatase activity was expressed as specific activity (nmol/min/mg) and as turnover number τ (s^-1^) each with the standard error of the fitted slope (SE). Activities were normalized to that of fully active Cn_WT_, which was defined as 100 %.

## RESULTS

### Calcineurin activation by Ca²⁺/calmodulin enhances NHE1-CnBD binding and stabilizes the complex

Binding of full-length heterodimeric human Cn (Cn_WT_) to NHE1 was quantified by fluorescence anisotropy (FA) using NHE1-CnBD*, a fluorescently labelled recombinant peptide comprising the NHE1 Cn-binding domain. For these measurements, Cn_WT_ and Cn_1-370_ were produced and purified reproducibly in three independent biological replicates (Fig. S1, Table S1). Dissociation constants (*K*_D_) were estimated by global fitting of all biological and technical replicates for each experimental condition (Fig. 1C-F). Confidence intervals were determined by profile likelihood analysis [36, 37] (Fig. 1G). The high quality of the datasets together with global parameter estimation yielded well-constrained *K*_D_ estimates with closed confidence intervals for all experimental conditions.

The constitutively active Cn_1-370_ was used for reference. It bound NHE1-CnBD* with a *K*_D_ of 7.9 [7.2; 8.7] nM (Fig. S2), consistent with previous data obtained by isothermal titration calorimetry with a slightly shorter NHE1-CnBD peptide (*K*_D_ 21 ± 2 nM) [7]. Cn_WT_ showed significantly lower affinity under the same Ca²⁺-containing conditions (*K*_D_ of 267 [249; 285] nM, Fig. 1D). At these conditions, Cn_WT_ is partially active (Fig. 1A, Table S1) [14]. Chelation of Ca^2+^ with EGTA, which completely inactivates Cn_WT_ (Fig. 1A, Table S1), did not alter binding affinity (*K*_D_ 260 [248; 271] nM, Fig. 1C, G). In contrast, addition of CaM in the presence of Ca^2+^ increased affinity approximately 7-fold (*K*_D_ 35 [33; 38] nM, Fig. 1E, G). Subsequent Ca^2+^ chelation in the CaM presence partially reversed this effect (*K*_D_ 148 [142; 153] nM, Fig. 1F, G) reflecting dissociation of CaM from Cn_WT_. The incomplete reversion can be explained by slow dissociation of CaM from Cn_WT_ after EGTA addition [14]. Together, these data show that Ca^2+^/CaM activation of full-length Cn markedly enhances NHE1-CnBD binding to tens of nanomolar affinity, but it is lower than that of constitutively active Cn_1-370_.

To assess the stability of the NHE1-CnBD−Cn complex, we analysed complexes of Cn_WT_ and NHE1-CnBD tagged with green fluorescent protein (GFP^NHE1-CnBD^) by quantitative analytical size-exclusion chromatography (qaSEC [26], Fig. 2). Cn_WT_ and GFP^NHE1-CnBD^ alone eluted at 12.9 ml and 14.6 ml, respectively (coloured dashed lines in Fig. 2A). Separation of an equimolar mixture of Cn_WT_ and GFP^NHE1-CnBD^ in the presence of Ca^2+^ resulted in elution of three partially overlapping peaks (Fig. 2A, top). The earliest peak (11.7 ml), contained nearly equimolar amounts of Cn_WT_ and GFP^NHE1-CnBD^, consistent with formation of a 1:1 complex. A second peak (12.6 ml) was enriched in Cn_WT_ (≈3.5 µM, concentration at peak maximum) but contained substantially less GFP^NHE1-CnBD^ (≈1 µM) (Fig. 2A) indicating partial dissociation of the complex during chromatography. The remaining free GFP^NHE1-CnBD^ eluted as third peak at 14.4 ml (Fig. 2A). Addition of CaM at equimolar concentration markedly altered the elution profile, with all three proteins co-eluting in a single peak at 11.3 ml (Fig. 2B). GFP^NHE1-CnBD^ and the Cn_WT_−CaM complex were present at nearly identical concentrations, consistent with a stable 1:1:1 complex. No noticeable dissociation of the complex was observed during qaSEC. As expected from the higher affinity, Cn_1-370_ formed a stable equimolar complex with GFP^NHE1-CnBD^ eluting as a single peak at 12.9 ml, which remained unchanged in the presence of CaM (Fig. 2C, Fig. S3).

**Fig. 2.**
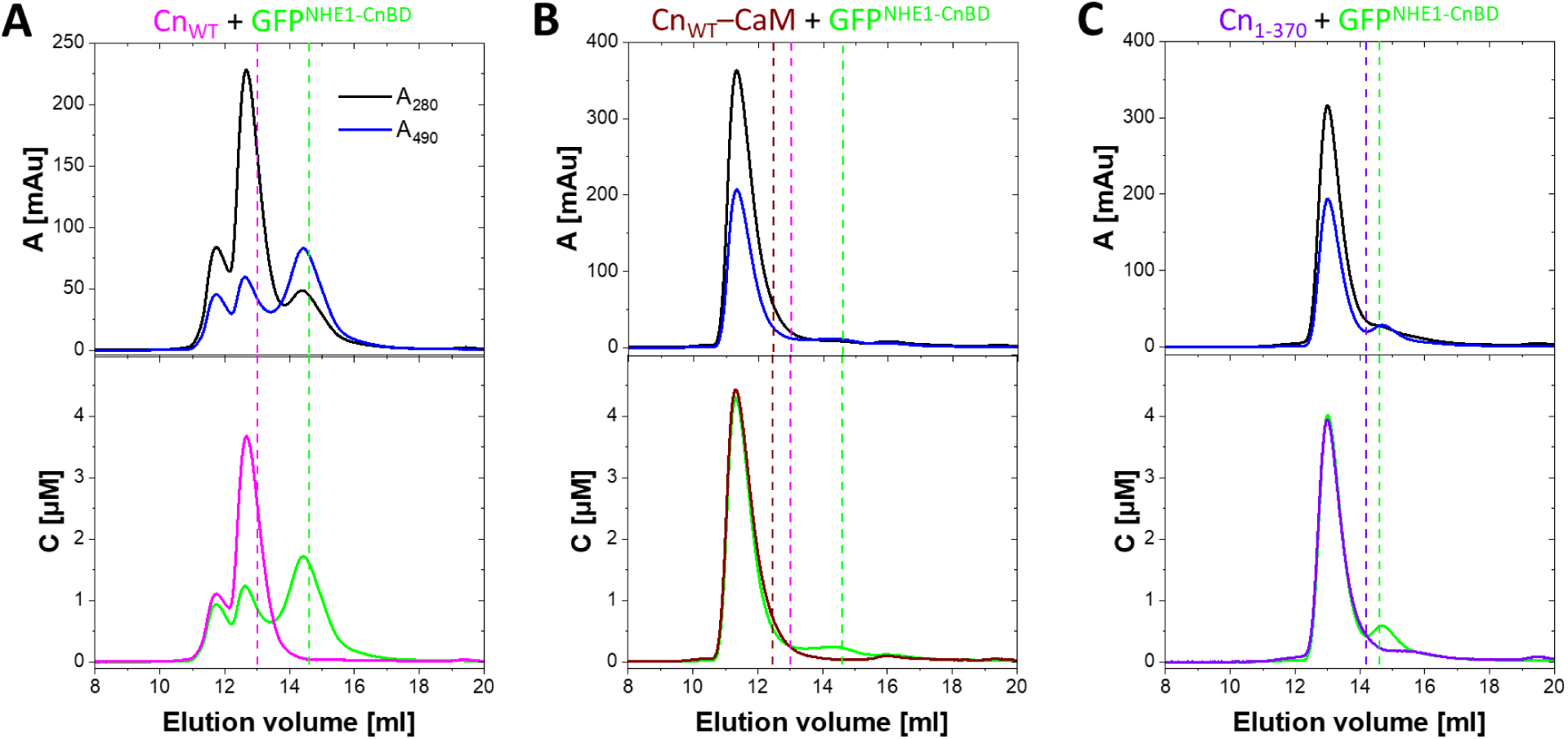
Activation of Cn_WT_ by Ca^2+^/CaM stabilizes its complex with NHE1-CnBD. Complex formation between calcineurin (Cn_WT_ or Cn_1-370_) and GFP-fused NHE1-CnBD (GFP^NHE1-CnBD^) was analysed by quantitative analytical size exclusion chromatography (qaSEC). All experiments were performed in the presence of Ca^2+^. Absorbance at 280 nm (A_280_, black) for total protein and at 490 nm (A_490_, blue) for GFP^NHE1-CnBD^ specifically was monitored (top panels) and used to quantify protein concentrations and complex stoichiometry across the elution profiles (bottom panels). Samples containing 10 µM of Cn_WT_ and GFP^NHE1-CnBD^ each (**A**); Cn_WT_, GFP^NHE1-CnBD^ and calmodulin (CaM) (**B**); or Cn_1-370_ and GFP^NHE1-CnBD^ (**C**) were analyzed, proteins are indicated with a respective colour above each panel. Elution maxima of the individual proteins analysed alone and of Cn_WT_–CaM complex are indicated by vertical dashed lines of the same colour. The nearly complete overlap of the Cn–CaM and Cn_1-370_ peaks with those of GFP^NHE1-CnBD^ (bottom panels **B** and **C**, respectively) indicates formation of stoichiometric complexes.

Together, the FA and qaSEC experiments demonstrate that Ca^2+^/CaM activation increases the affinity of full-length Cn for NHE1-CnBD and stabilizes the resulting complex.

### NHE1-CnBD competes with the cyclophilin A−cyclosporin A complex for calcineurin binding

NHE1 was shown to bind to both the LxVP- and PxIxIT-binding sites of Cn, which are critical for substrate recognition [7, 13]. As the inhibitory CsA/CypA complex binds as well to the LxVP-binding site [14, 19], we analysed whether NHE1-CnBD competes with the CsA/CypA complex for Cn binding. We quantified the effect of the CsA/CypA complex on the Cn-NHE1 interaction using FA (Fig. 3A-D). In the presence of 15 µM CsA/CypA, NHE1-CnBD binding to Cn_WT_ was markedly reduced, with the estimated apparent dissociation constant (*K*_D_ ^app^) increasing to 2.96 [2.69; 3.26] µM (Fig. 3A). A similar competition effect was observed with Cn_WT_ in the presence of Ca^2+^/CaM, yielding a *K*_D_ ^app^ of 1.44 [1.34; 1.53] µM (Fig. 3B). In contrast, CypA alone had only a minor effect on the interaction, increasing the *K*_D_ ^app^ to 361 [345; 377] nM (Fig. 3C), slightly above the value measured in the absence of additives (Fig. 3D).

**Fig. 3.**
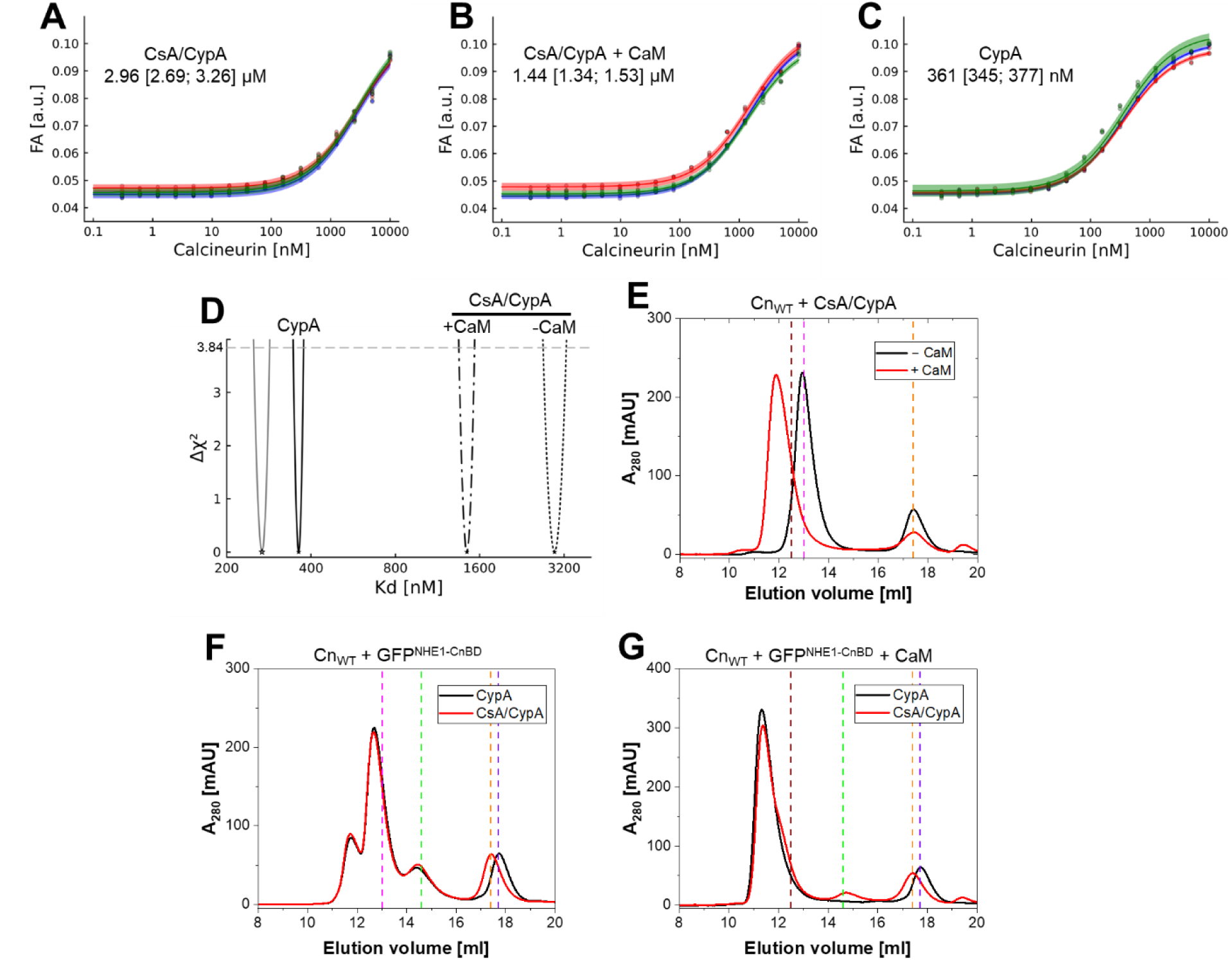
NHE1-CnBD outcompetes CsA/CypA binding to Cn_WT_. **A-C**, Binding (*K*_D_^app^) of full-length Cn_WT_ to the fluorescein-labeled NHE1-CnBD peptide was quantified with FA with addition of CsA/CypA in the absence (A) or presence (B) of 10 µM CaM, and with addition of CypA only (**C**). All measurements were performed in the FA buffer containing 1 mM Ca^2+^ with three independent biological replicates of Cn_WT_ (in different colour), each with three technical replicates. The combined data from individual experiments were fitted with a 1:1 binding model using global non-linear regression. Determined *K*_D_^app^ values with respective CI’s are shown in each panel. **D**, 95 % CI’s (Δχ^2^ of 3.84) of *K*_D_ values for measurements shown in (**A-C**) calculated with profile likelihood method. The CI of *K*_D_ value for Cn_WT_/NHE1 CnBD interaction without any addition is shown in grey for comparison (the first, unlabelled curve). **E**, CsA/CypA forms a stable complex with Cn_WT_ only in the presence of Ca^2+^/CaM, but not in the absence of CaM, in analytical SEC. **F-G**, The SEC elution profiles of Cn_WT_/GFP^NHE1-CnBD^ are not affected by CsA/CypA addition in the absence (**F**) of CaM, whereas and CsA/CypA show only minor effect in the CaM presence (**G**). In **F-G**, samples were analysed either in the presence of CypA only (black lines) or the CsA/CypA complex (red lines). Vertical dashed lines in **E-G** show positions of the elution maxima of reference proteins and complexes (magenta, Cn_WT_; green, GFP^NHE1-CnBD^; orange, CsA/CypA complex; brown, Cn_WT_–CaM complex; violet, CypA). Cn_WT_, GFP^NHE1-CnBD^, and CaM were used at 10 µM in analytical SEC; CypA and CsA were used at 15 µM both in FA assay and in analytical SEC.

To further examine the effect of CsA/CypA on Cn−NHE1-CnBD complex formation, we analyzed the interaction by analytical SEC. CsA/CypA addition resulted in reduction of the retention volume of Cn_WT_ in the presence of Ca^2+^/CaM, but not without CaM (Fig. 3E), indicating formation of a stable inhibited complex only in the presence of CaM under gel filtration condition. Further, CsA/CypA addition did not substantially alter the elution profile of the Cn_WT_−GFP^NHE1-CnBD^ complex as compared to that of CypA addition alone (Fig. 3F) as well as to that of the GFP^NHE1-CnBD^ and Cn_WT_ without inhibitory components (Fig. 2A). In the presence of CaM, addition of CsA/CypA had only a minimal effect on the Cn_WT_−GFP^NHE1-CnBD^ complex (Fig. 3G). At most, a slight reduction in the Cn_WT_−CaM−GFP^NHE1-CnBD^ peak was accompanied by the appearance of a small shoulder on its trailing edge, most likely corresponding to the Cn_WT_−CaM−CsA/CypA complex. Consistent with this interpretation, a small peak appeared at 14.7 ml, corresponding to free GFP^NHE1-CnBD^ displaced from the complex. Thus, under the conditions of analytical SEC, CsA/CypA had only a minor effect on the Cn_WT_−NHE1-CnBD.

Together, these results indicate that NHE1-CnBD and the CsA/CypA complex compete for an overlapping binding site on Cn, consistent with engagement of the LxVP-binding site. Moreover, the NHE1-CnBD binding to Cn appears to be stronger than that of the commonly used inhibitor complex CsA/CypA, as it could not replace NHE1-CnBD under gel filtration conditions.

### Binding of NHE1-CnBD inhibits the phosphatase activity of calcineurin towards protein substrate

As NHE1-CnBD binds Cn with high affinity and competes with the inhibitory CsA/CypA complex for the LxVP-binding site, we next examined the effect of NHE1-CnBD binding on Cn phosphatase activity. Firstly, the RII phosphopeptide, a standard peptide substrate of Cn, was used to quantify the phosphatase activity. In the presence of Ca^2+^/CaM, Cn_WT_ showed a specific activity of 480 ± 10 nmol min^-1^ mg^-1^, approximately 40-fold higher than that of Cn_WT_ in the absence of CaM (12 ± 9 nmol min^-1^ mg^-1^) (Fig. 4A, Table S2). Addition of NHE1-CnBD in the presence of CaM inhibited RII dephosphorylation in a concentration-dependent manner. At 0.05 µM NHE1-CnBD, Cn_WT_ activity decreased to 68 % of the control, and was further reduced to 26 % at 0.1 µM. At 1 µM NHE1-CnBD, phosphatase activity was almost completely suppressed, reaching a level comparable to that of Cn_WT_ in the absence of CaM (Fig. 4A, Table S2).

**Fig. 4.**
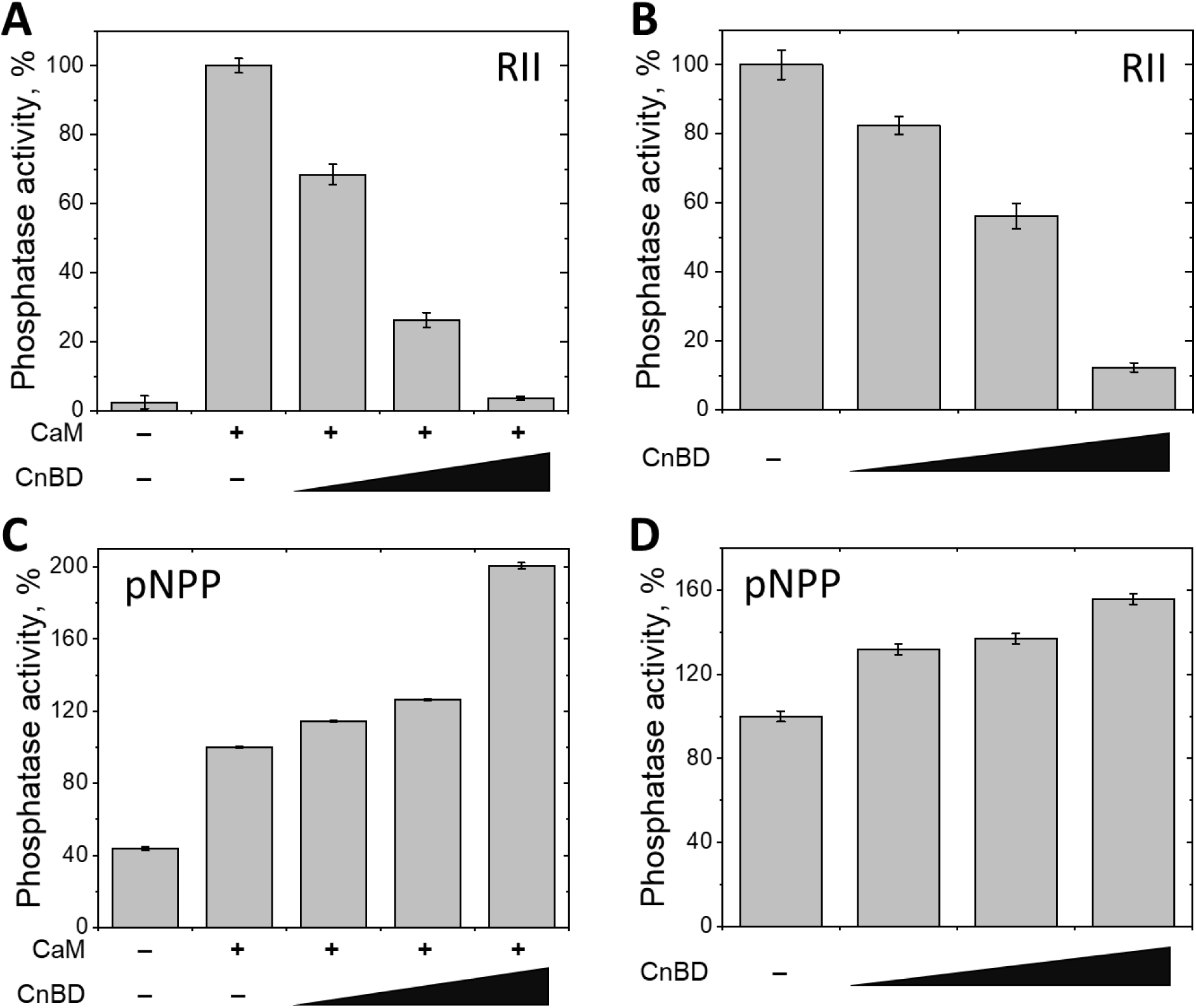
NHE1-CnBD inhibits the specific Cn phosphatase activity and increases its non- specific activity. The Cn phosphatase activity was measured using its specific model substrate RII phosphopeptide (**A, B**) or non-specific phosphatase substrate pNPP (**C, D**) for the full- length Cn_WT_ (**A, C**) or constitutively active Cn_1-370_ (**B, D**) in the absence or with addition of 0.05 µM, 0.1 µM, or 1.0 µM NHE1-CnBD. Cn activity in the fully active state was taken as 100 %, activity in the partially active state is shown for comparison in **A** and **C**. Measurements were performed in triplicate. Error bars represent the standard errors of the linear fit to the initial part of the reaction curve.

A similar inhibitory effect was observed for Cn_1-370_ (Fig. 4B). Although it showed approximately fourfold higher specific activity than Cn_WT_ in the presence of Ca^2+^/CaM (Table S2), RII dephosphorylation activity was reduced to 82 %, 60 %, and 13 % of the control in the presence of 0.05 µM, 0.1 µM, and 1.0 µM NHE1-CnBD, respectively (Fig. 4B, Table S2).

The effect was markedly different when pNPP, a small-molecule substrate requiring access only to the catalytic site, was used. In presence of Ca^2+^/CaM, NHE1-CnBD enhanced phosphatase activity in a concentration dependent manner. At 0.05 µM and 0.1 µM NHE1-CnBD, pNPP dephosphorylation by Cn_WT_ was increased by 15 % and 26 %, respectively, whereas 1.0 µM NHE1-CnBD increased activity approximately 100 % (Figure 4C, Table S3). In the absence of CaM, pNPP dephosphorylation by Cn_WT_ was about 40 % of the Cn_WT_ activity in presence of Ca^2+^/CaM (Fig. 4C); and NHE1-CnBD substantially increased pNPP dephosphorylation rate only at 1.0 µM (Fig. S4, Table S3), consistent with a lower affinity of partially active Cn_WT_. NHE1-CnBD also stimulated Cn_1-370_ pNPP phosphatase activity in a concentration-dependent manner, yet at 1 µM NHE1-CnBD, the activity only increased by 55 % (Figure 4D, Table S3).

Together, these results demonstrate that NHE1-CnBD binds full-length Cn with high affinity, and that Ca^2+^/CaM-dependent activation of Cn further strengthens this interaction by more than sevenfold, increasing the affinity from the submicromolar to the tens of nanomolar range. High-affinity binding of NHE1-CnBD efficiently suppresses dephosphorylation of a model peptide substrate while leaving the catalytic site intact, as shown by the stimulation of pNPP hydrolysis. It therefore acts as a typical Cn inhibitor, even outcompeting the inhibitory CsA/CypA complex for Cn binding. These findings indicate that NHE1-CnBD acts as a substrate-competitive inhibitor of Cn by occupying the LxVP-binding site and preventing substrate recognition.

## DISCUSSION

The present study extends previous analyses of the NHE1−Cn interaction by examining full-length heterodimeric Cn in distinct activation states. Whereas enhanced ligand binding upon Ca^2+^/CaM activation of Cn is commonly accepted based on qualitative interaction studies and structural characterization [13, 14, 16, 17], its quantitative effect on the affinity of full-length Cn for NHE1 has not previously been determined. As expected, Ca^2+^/CaM activation significantly increased the affinity of Cn for NHE1-CnBD, consistent with greater accessibility of the LxVP-binding site. Nevertheless, the submicromolar affinity observed for both the inactive and partially active states remained considerably higher than previously reported for NHE1-CTD (residues 680-815) lacking the LTVP motif [7]. This finding suggests that the LxVP-binding site is already at least partially accessible before full activation, allowing productive binding of the LTVP motif. This interpretation is consistent with structural models proposing that the regulatory region only partially occludes the LxVP-binding site and that Ca^2+^/CaM-dependent displacement of the autoinhibitory domain (AID) fully exposes this interface [25]. Furthermore, the lower affinity of fully active full-length Cn (*K*_D_ of 35 [33; 38] nM, Fig. 1E) compared with the constitutively active Cn_1-370_ (7.9 [7.2; 8.7] nM, Fig. S2) suggests that the regulatory region and the AID continue to influence ligand binding even after activation.

Our findings also extend previous observations that Ca^2+^/CaM activation promotes Cn interactions with diverse binding partners. Enhanced binding of NFAT1 and Ca_V_3.2 to Ca^2+^/CaM-activated Cn has previously been demonstrated using pull-down assays [16, 17], but these studies neither quantified binding affinities nor addressed the stability of the resulting complexes. The qaSEC analysis presented here complements these observations and our affinity measurements by demonstrating that the stability of the NHE1-CnBD−Cn complex is also regulated by the activation state of Cn. In the partially active state, Cn formed a heterogenous mixture of stable and dissociating NHE1-CnBD−Cn complexes present along with free protein, consistent with a dynamic binding equilibrium. In contrast, Ca^2+^/CaM- activation markedly stabilized the complex, in agreement with the increased binding affinity determined by FA. These findings indicate that Ca^2+^/CaM-dependent activation of Cn regulates not only equilibrium affinity but also the stability of Cn-substrate or regulator complexes. Such effects on complex lifetime could represent a general mechanism by which Ca^2+^ signals tune substrate recognition and signalling by Cn. In the case of NHE1, activation-dependent stabilization of the complex is expected to facilitate the reciprocal regulation of both proteins [7, 8, 13], promoting sustained association under conditions of activation while permitting more dynamic interactions at lower activation states.

Previous work showed that an NHE1-derived peptide containing only its PxIxIT-like motif (PIVITID) does not affect Cn phosphatase activity [8]. Subsequent studies demonstrated that inclusion of the upstream LTVP motif enables bipartite binding to constitutively active truncated Cn with nanomolar affinity through avidity [7]. Here, we extend these findings to full-length Cn and show that binding with nanomolar affinity is retained despite activation-dependent regulation of the LxVP-binding site. Affinities in this range are characteristic of endogenous Cn regulators and inhibitors, including CABIN1 and RCAN1, scaffold proteins, viral inhibitors such as A238L, and the CsA/CypA and FK506/FKBP12 inhibitory complexes [22–24, 38–40] [41], [42], whereas physiological substrates typically bind with micromolar affinity [15, 43]. Like the clinically used immunosuppressants CsA and FK506 immunophilin complexes [44], NHE1 occupies the LxVP-binding site and prevents substrate recognition. Among known Cn ligands, the African swine fever virus inhibitor A238L, most closely resembles the binding mode of NHE1 by simultaneously engaging the PxIxIT- and LxVP-binding sites through a bipartite interaction, binding constitutively active Cn with a *K*_D_ of ∼4 nM [36]. High-affinity multivalent binding is generally incompatible with rapid substrate turnover and instead favours the formation of stable regulatory complexes. Consistent with this consideration, Thr779 within NHE1 is dephosphorylated by constitutively active Cn only very slowly [7], despite residing in suitable distance downstream of the Cn-binding region in the NHE1-CTD. Moreover, NHE1-CnBD potently inhibited dephosphorylation of the RII phosphopeptide substrate at nanomolar concentrations while simultaneously enhancing hydrolysis of the small-molecule substrate pNPP. Together with the competition experiments using the CsA/CypA complex, these finding indicate that NHE1 inhibits Cn primarily by blocking substrate recruitment rather than by impairing the catalytic site. Thus, despite similarities in their multivalent binding strategies, NHE1 appears mechanistically more similar to the viral inhibitor A238L than to RCAN1 and binds more efficiently than the CsA/CypA complex.

The physiological consequences of high-affinity NHE1-Cn binding remain to be established. The potent inhibition of Cn by NHE1-CnBD appears difficult to reconcile with reports that NHE1 overexpression promotes Cn/NFAT signalling in cardiomyocytes in an NHE1 activity-dependent manner [8]. One possible explanation is that NHE1 primarily functions as a membrane-localized scaffold that recruits Cn to the plasma membrane. Given that Cn and NHE1 are ubiquitously expressed but that Cn is considerably more abundant than NHE1 in most tissues, only a fraction of the cellular phosphatase would be expected to be membrane associated at any time. Dynamic association with NHE1 could therefore increase the local concentration of Cn without globally activating the enzyme at basal cellular Ca^2+^ concentrations. Our data further suggest that NHE1-bound Cn is maintained in a substrate-competitive inhibited state. Changes in intracellular Ca^2+^/CaM concentration, CaM occupancy or local physicochemical conditions such as pH may shift this equilibrium, allowing Cn to dissociate and act on nearby substrates.

In summary, our results provide a quantitative analysis of activation-dependent binding of full-length Cn and identify NHE1 as a high-affinity, membrane associated regulator of Cn that combines scaffold and substrate-competitive inhibitory functions, rather than serving solely as a docking site for substrate dephosphorylation. In this respect, NHE1 resembles the recently described small membrane protein calcimembrin (C16orf74), which likewise binds Cn with high affinity and suppresses phosphatase activity [21]. Beyond providing mechanistic insight into reciprocal regulation of NHE1 and Cn, the quantitative data reported here may also facilitate the development of NHE1-inspired bivalent Cn ligands as selective modulators of Cn signalling.

## ABBREVIATIONS

AID: autoinhibitory domain
CABIN1: calcineurin-binding protein 1
CaM: calmodulin
Cn: calcineurin
CsA: cyclosporin A
CypA: cyclophilin A
EGTA: ethylene glycol-bis(β-aminoethyl ether)-N,N,N′,N′-tetraacetic acid
FA: fluorescence anisotropy
FKBP12: FK506-binding protein 12
GFP^NHE1-CnBD^: fusion construct of green fluorescent protein and NHE1-CnBD
IPTG: isopropyl-β-D-1-thiogalactopyranoside
NFAT: nuclear factor of activated T-cell
NHE1: Na^+^/H^+^-exchanger 1
NHE1-CnBD: NHE1 calcineurin-binding domain (residues 674-730)
NHE1-CTD: NHE1 C-terminal domain
pNPP: *para*-nitrophenyl phosphate
qaSEC: quantitative analytical size-exclusion chromatograpy
RCAN1: regulator of calcineurin 1
RII: regulatory subunit type II of protein kinase A
SLiM: short linear motif
TCEP: tris(2-carboxyethyl)phosphine
TEV: tobacco etch virus

## ACKNOWLEDGEMENTS

This work was supported by the Deutsche Forschungsgemeinschaft (DFG, German Research Foundation) under Germany’s Excellence Strategy (CIBSS – EXC-2189 – Project ID 390939984) in the form of project funding to C.H. and V.T., and of CIBSS Launchpad Program Funds to E.V.M.; by the DFG through Project-ID 431984000 – SFB 1453 to C.H. and through Project-ID 403222702 – SFB 1381 to C.H. and V.T.; by the Baden-Württemberg Stiftung through Project ‘ArrhythMEC’ to V.T.; European Union’s Horizon Europe Research and Innovation Programme under Grant Agreement No. 101136299 project ARTEMIs (AcceleRating the Translation of Virtual Twins towards a Personalised Management of Steatotic Liver Patients) to M.H. We thank Lukas Kmetsch and Christophe Wirth for critical discussion of the manuscript.

## AUTHOR CONTRIBUTIONS

CH and EVM conceived and designed the study. AS, JSH, SG, BAK, and EVM performed experiments. AS, MH, VT, EVM, and CH analysed the data. EVM and CH drafted the manuscript. All authors reviewed and approved the final version of the manuscript.

## AI DISCLOSURE

The authors used ChatGPT (OpenAI, San Francisco, CA, USA) and DeepL Write (DeepL SE, Cologne, Germany) for language editing. The authors reviewed and edited the output and take full responsibility for the content of the publication.

## Notes

### Competing Interest Statement

The authors have declared no competing interest.

