## Supplementary for "Human NHE1 binds full-length calcineurin with high affinity and inhibits substrate dephosphorylation"

1 – Institute of Biochemistry and Molecular Biology, ZBMZ, Faculty of Medicine, University of Freiburg, Freiburg, Germany; 2 – Department of Life Sciences, University of Applied Sciences Albstadt-Sigmaringen, Sigmaringen, Germany; 3 – Freiburg Center for Data Analysis and Modelling (FDM), University of Freiburg, Freiburg, Germany; 4 – CIBSS - Centre for Integrative Biological Signalling Studies, University of Freiburg, Freiburg, Germany; 5 – Institute of Physics, University of Freiburg, Freiburg, Germany; 6 – Institute of Molecular Medicine and Cell Research, University of Freiburg, Freiburg, Germany

*contributed equally, #correspondence to: Institute of Biochemistry and Molecular Biology, ZBMZ, Faculty of Medicine, CIBSS, University of Freiburg, Stefan-Meier-Strasse 17, 79104 Freiburg, Germany; (EVM), (CH)

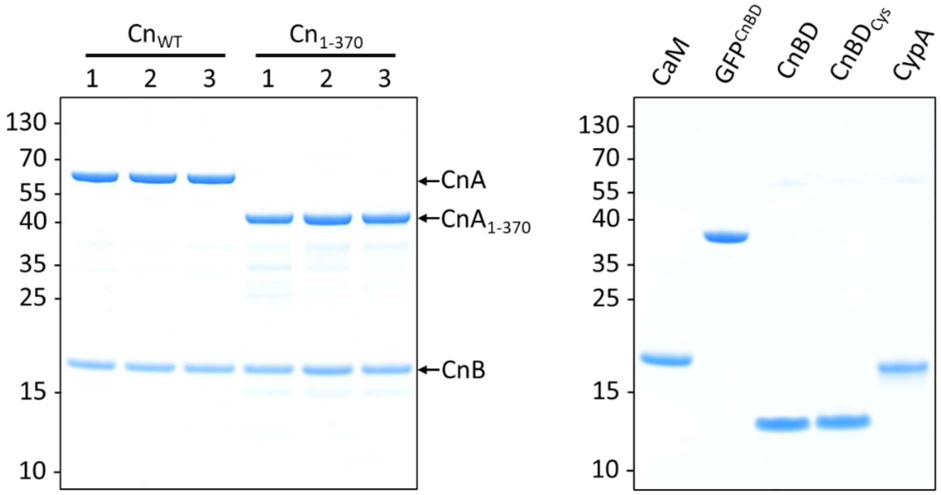


**Fig. S1. Quality control of three biological replicates of full-length calcineurin ($Cn_{WT}$), constitutively active calcineurin ($Cn_{1\text{-}370}$) and other proteins used in the study.** Purified proteins were analyzed by SDS-PAGE using 4-12 % NuPAGE Bis-tris gels. Positions of calcineurin A (CnA) and B (CnB) are indicated in the left gel. Positions of molecular mass standard proteins (kDa) are shown on the left side of each gel. CaM: calmodulin, CnBD: NHE1 calcineurin-binding domain (NHE1-CnBD, residues 674-730), $CnBD_{Cys}$: NHE1-CnBD with S729C point mutation, $GFP^{CnBD}$: fusion construct of GFP and NHE1-CnBD ($GFP^{NHE1\text{-}CnBD}$), CypA: cyclophilin A.

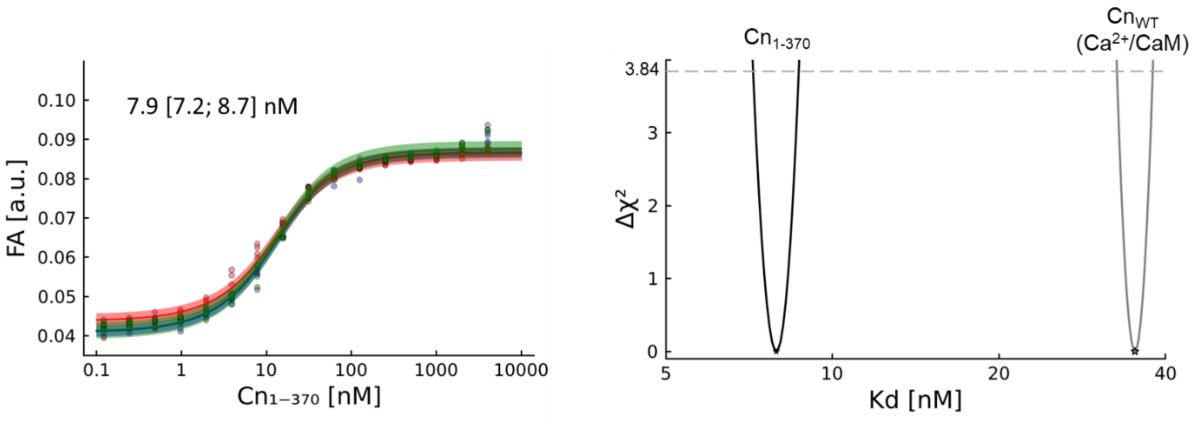


**Fig. S2. Affinity ($K_D$) of $Cn_{1\text{-}370}$ for NHE1-CnBD* determined with FA assay.** Measurements were performed in the presence of $Ca^{2+}$ with three independent biological replicates, each with four technical replicates. Fitting of combined data (left) and CI calculation (right) were done as described for other FA assays (see Figs. 1 and 3). CI for $Cn_{WT}$ in the presence of $Ca^{2+}$/CaM is shown with a grey curve in the right plot for comparison.

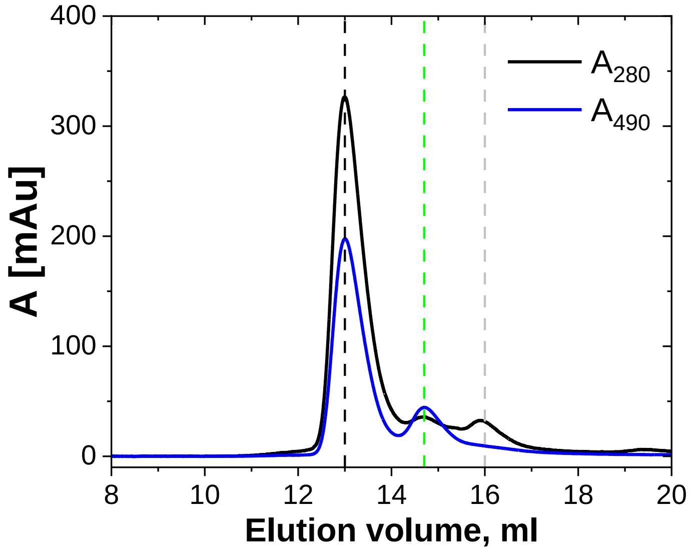


**Fig. S3. aSEC of the sample containing $Cn_{1-370}$, $GFP^{NHE1-CnBD}$ and CaM.** $A_{280}$ for total protein and $A_{490}$ for $GFP^{NHE1-CnBD}$ were monitored. For reference, positions of elution maxima of $Cn_{1-370}$–$GFP^{NHE1-CnBD}$ complex (see. Fig. 2C), $GFP^{NHE1-CnBD}$, and CaM are indicated with black, green and grey dashed lines, respectively. Note the same position and height of the peak corresponding to the $Cn_{1-370}$–$GFP^{NHE1-CnBD}$ complex as in the absence of CaM. In this sample, CaM elutes separately from the $Cn_{1-370}$–$GFP^{NHE1-CnBD}$ complex at about 16.0 ml.

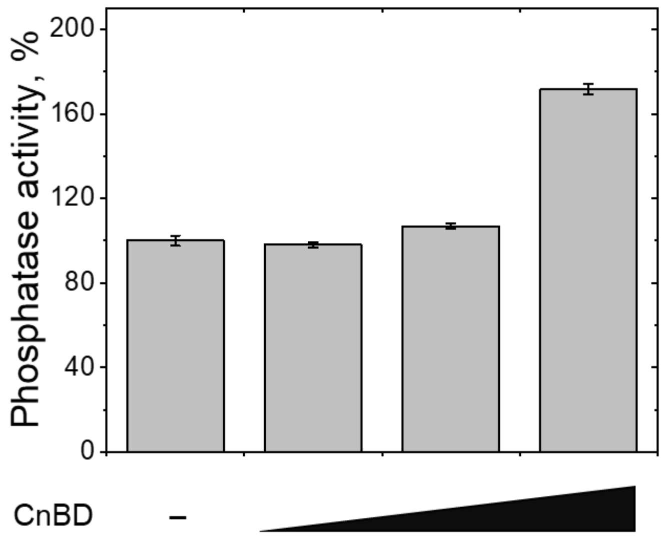


**Fig. S4. NHE1-CnBD enhances pNPP dephosphorylation by partially active $Cn_{WT}$.** pNPP dephosphorylation rate was measured in the absence (set as 100 %) or with addition of 0.05 µM, 0.1 µM, or 1.0 µM NHE1-CnBD**.**

**Table S1. pNPP dephosphorylation activity of three batches (biological replicates) of full-length calcineurin ($Cn_{WT}$) and activated variant ($Cn_{1\text{-}370}$) in different activation states.**

The activity was measured in the assay buffer containing 1 mM $CaCl_2$ with addition, where mentioned, of 5 mM EGTA or 0.1 µM calmodulin (CaM). Measured *para*-nitrophenol release (v) for $Cn_{WT}$ and $Cn_{1\text{-}370}$ in the presence of EGTA is the range of a technical error meaning no detectable enzymatic activity (*).

| batch | state | additive | Cn [nM] | v[a] [nmol/min] | specific activity [nmol/min/mg] | τ [$s^{-1}$] |
|---|---|---|---|---|---|---|
| **blank** | | EGTA | 0 | 0.007 ± 0.000 | - | - |
| | | - | 0 | 0.004 ± 0.004 | - | - |
| | | CaM | 0 | 0.007 ± 0.000 | - | - |
| **$Cn_{WT}$ 1** | inactive | EGTA | 25 | 0.002 ± 0.000 | 11 ± 0* | 0.01 ± 0.00* |
| | part. active | - | 50 | 0.368 ± 0.013 | 925 ± 34 | 1.23 ± 0.04 |
| | fully active | CaM | 25 | 0.382 ± 0.011 | 1918 ± 56 | 2.54 ± 0.07 |
| **$Cn_{WT}$ 2** | inactive | EGTA | 25 | 0 | 0* | 0* |
| | part. active | - | 50 | 0.364 ± 0.009 | 914 ± 22 | 1.21 ± 0.03 |
| | fully active | CaM | 25 | 0.415 ± 0.002 | 2086 ± 11 | 2.77 ± 0.01 |
| **$Cn_{WT}$ 3** | inactive | EGTA | 25 | 0.002 ± 0.000 | 11 ± 0* | ≈ 0* |
| | part. active | - | 50 | 0.435 ± 0.007 | 1094 ± 17 | 1.45 ± 0.02 |
| | fully active | CaM | 25 | 0.449 ± 0.002 | 2255 ± 11 | 2.99 ± 0.01 |
| **$Cn_{1\text{-}370}$ 1** | inactive | EGTA | 10 | 0.004 ± 0.000 | 56 ± 0* | 0.07 ± 0.00* |
| | active | - | 10 | 0.087 ± 0.004 | 1094 ± 56 | 1.45 ± 0.07 |
| **$Cn_{1\text{-}370}$ 2** | inactive | EGTA | 10 | 0.004 ± 0.002 | 56 ± 28* | 0.07 ± 0.04* |
| | active | - | 10 | 0.176 ± 0.004 | 2215 ± 56 | 2.94 ± 0.07 |
| **$Cn_{1\text{-}370}$ 3** | inactive | EGTA | 10 | 0.002 ± 0.002 | 28 ± 28* | 0.04 ± 0.04* |
| | active | - | 10 | 0.161 ± 0.004 | 2019 ± 56 | 2.68 ± 0.07 |

[a]Values are slopes of the initial linear part of the reaction curve with standard errors of a linear fit.

**Table S2. NHE1 CnBD inhibits a specific calcineurin phosphatase activity.**

Dephosphorylation of RII phosphopeptide was measured in $Ca^{2+}$-containing activity buffer in the presence of 0-1.0 µM NHE1 CnBD peptide. Biological replicate #3 of $Cn_{WT}$ and #2 of $Cn_{1-370}$ at 10 nM were used.

| | CaM [µM] | NHE1-CnBD [µM] | Specific activity [nmol/min/mg] | τ [$s^{-1}$] |
|---|---|---|---|---|
| **$Cn_{WT}$** | 0 | 0 | 12 ± 9 | 0.197 ± 0.012 |
| | 0.1 | 0 | 480 ±10 | 7.996 ± 0.013 |
| | 0.1 | 0.05 | 328 ± 1 | 5.472 ± 0.002 |
| | 0.1 | 0.1 | 126 ± 10 | 2.103 ±0.013 |
| | 0.1 | 1.0 | 17 ± 3 | 0.285 ± 0.004 |
| **$Cn_{1-370}$** | 0 | 0 | 1888 ± 86 | 31.46 ± 0.09 |
| | 0 | 0.05 | 1554 ± 53 | 25.90 ± 0.06 |
| | 0 | 0.1 | 1126 ± 74 | 18.82 ± 0.08 |
| | 0 | 1.0 | 247 ± 26 | 4.12 ± 0.03 |

**Table S3. pNPP phosphatase activity of calcineurin is increased by NHE1-CnBD.**

Biological replicate #2 of $Cn_{WT}$ and #2 of $Cn_{1-370}$ were used, respective activity in the absence of NHE1-CnBD is shown in Table S1.

| Sample | CaM [µM] | NHE1-CnBD [µM] | Cn [nM] | V [nmol/min] | specific activity [nmol/min/mg] | τ [$s^{-1}$] |
|---|---|---|---|---|---|---|
| **blank** | 0 | 0.05 | 0 | 0.011 ± 0.002 | - | - |
| | 0 | 0.1 | 0 | 0.013 ± 0.002 | - | - |
| | 0 | 1.0 | 0 | 0.013 ± 0.000 | - | - |
| **$Cn_{WT}$** | 0 | 0.05 | 25 | 0.179 ± 0.002 | 897 ± 11 | 1.19 ± 0.01 |
| | 0 | 0.1 | 25 | 0.194 ± 0.002 | 976 ± 11 | 1.29 ± 0.01 |
| | 0 | 1.0 | 25 | 0.313 ± 0.004 | 1570 ± 22 | 2.08 ± 0.03 |
| | 0.1 | 0.05 | 25 | 0.475 ± 0.002 | 2389 ± 11 | 3.17 ± 0.01 |
| | 0.1 | 0.1 | 25 | 0.525 ± 0.002 | 2636 ± 11 | 3.50 ± 0.01 |
| | 0.1 | 1.0 | 25 | 0.833 ± 0.007 | 4184 ± 34 | 5.55 ± 0.04 |
| **$Cn_{1-370}$** | 0 | 0.05 | 10 | 0.232 ± 0.004 | 2916 ± 56 | 3.87 ± 0.07 |
| | 0 | 0.1 | 10 | 0.241 ± 0.004 | 3029 ± 56 | 4.02 ± 0.07 |
| | 0 | 1.0 | 10 | 0.275 ± 0.004 | 3449 ± 56 | 4.58 ± 0.07 |
